# The Coding And Small-Non-Coding Hippocampal Synaptic RNAome

**DOI:** 10.1101/2020.11.27.401901

**Authors:** Robert Epple, Dennis Krüger, Tea Berulava, Gerrit Brehm, Rezaul Islam, Sarah Köster, Andre Fischer

## Abstract

Neurons are highly compartmentalized cells that depend on local protein synthesis. Thus, messenger RNAs (mRNAs) have been detected in neuronal dendrites and more recently also at the pre- and postsynaptic compartment. Other RNA species, such as microRNAs, have also been described at synapses where they are believed to control mRNA availability for local translation. Nevertheless, a combined dataset analyzing the synaptic coding and non-coding RNAome via next-generation sequencing approaches is missing. Here we isolate synaptosomes from the hippocampus of young wild type mice and provide the coding and non-coding synaptic RNAome. These data are complemented by a novel approach to analyze the synaptic RNAome from primary hippocampal neurons grown in microfluidic chambers. Our data show that synaptic microRNAs control almost the entire synaptic mRNAome and we identified several hub microRNAs. By combining the *in vivo* synaptosomal data with our novel microfluidic chamber system, we also provide evidence to support the hypothesis that part of the synaptic microRNAome may be supplied to neurons via astrocytes. Moreover, the microfluidic system is suitable to study the dynamics of the synaptic RNAome in response to stimulation. In conclusion, our data provide a <u>valuable</u> resource and hint to several important targets for future experiments.

## Introduction

Neurons are highly compartmentalized cells that form chemical synapses and the plasticity of such synapses is a key process underlying cognitive function. In turn, loss of synaptic integrity and plasticity is an early event in neuropsychiatric and neurodegenerative diseases. Synapses are usually far away from the soma, which raises the question how neurons ensure the supply of synaptic proteins. Theoretical considerations and a substantial amount of data show that mRNAs coding for key synaptic proteins are transported along dendrites to synaptic compartments, where they are locally translated into proteins (Doyle & Kiebler, 2011) (Kosik, 2016) (Holt *et al*, 2019) (Fonkeu *et al*, 2019) (Biever, 2020). Hence, several studies investigated the synaptic RNAome via different approaches. For example, early *in situ* hybridization experiments demonstrated the localization of specific mRNAs to synapses (Garner *et al*, 1988). In addition, microarray and RNA-seq techniques were used to study the synapto-dendritic (Cajigas *et al*, 2012) (Ainsley *et al*, 2014) (Farris *et al*, 2019), synapto-neurosomal (Most *et al*, 2015) and more recently also the synaptosomal RNA pool of the mouse brain (Chen *et al*, 2017) (Hafner, 2019). However, compared to mRNAs, there is comparatively less knowledge about the non-coding RNAome at synapses. The best known non-coding RNAs are microRNAs which are 19-22 nucleotide long RNA molecules regulating protein homeostasis via binding to a target mRNA thereby causing its degradation or inhibition of translation (Gurtan & Sharp, 2013). Several microRNAs have been implicated with synaptic plasticity and were identified at synapses where they have been linked to the regulation of mRNA stability and availability for translation (Smalheiser, 2014) (Weiss *et al*, 2015) (Rajman & Schratt, 2017) (Sambandan *et al*, 2017). The combined analysis of the synaptic microRNA/mRNAome is however lacking and knowledge about other non-coding RNA species is rare. Another issue is that the methods used so far to study synaptic RNAs from tissue samples do not allow to distinguish between RNAs produced by the corresponding neurons and RNAs that might be transferred to synapses from other cell types. This question is becoming increasingly important, since there is emerging evidence for inter-cellular RNA transport and data supporting the hypothesis that for example glia cells provide neurons with RNA (Sotelo *et al*, 2014) (Jose, 2015). In this study we isolated synaptosomes from the hippocampus of mice and performed from the same preparation total and smallRNA-sequencing. To complement these data and address the question about the origin of synaptic RNAs we developed a novel microfluid chamber that not only allowed us to grow primary hippocampal neurons that form synapses in a pre-defined compartment (Taylor *et al*, 2010), but enabled us to isolate the synaptic compartments from these chambers using a novel device we call SNIDER (SyNapse Isolation DevicE by Refined Cutting) followed by RNA-sequencing. We also show that this novel microfluid chamber is suitable to assay the dynamics of the synaptic RNAome in response to stimulation. In conclusion, our experiments allowed us for the first time to build a high-quality synaptic microRNA/mRNA network and suggest key synaptic RNAs, including lncRNAs and snoRNAs, for future mechanistic studies in the context of the healthy and diseased brain.

## Results

### The <u>hippocampal</u> coding and non-coding <u>synaptosomal</u> RNAome

<u>We isolated high-quality synaptosomes from the hippocampus of 3 months old mice, and processed the corresponding RNA for total and small RNA-sequencing</u> **(Fig 1A).** After quality control for high confidence transcripts we could detect 234 mRNA, 6 lncRNAs (excluding sequences that code for predicted genes), 65 microRNAs and 37 SnoRNAs **(Fig 1B, tables S1, 2, and 3).** GO-term analysis revealed that the mRNAs reflect exclusively the pre-and post-synaptic compartment **(Fig 1C)** confirming the quality of our data. Functional pathway analysis showed that the mRNAs found in our synaptosomal preparations represent key pathways linked to synaptic function and plasticity **(Fig. 1D).** We also observed a substantial amount of highly abundant microRNAs present in our synaptosomal preparations **(Table S2)** <u>and</u> wanted to understand the synaptic regulatory mRNA-microRNA network. To this end, we applied a novel bioinformatic approach and first generated the mRNA-network using the mRNAs detected at synapses, intersected this network with the synaptic microRNAome and asked if any of mRNAs within the network represent confirmed microRNA targets. Our data revealed that the 98% of the synaptic mRNAome is targeted by 95% of the synaptic microRNAs **(Fig 1E, F).** These data suggest that the synaptic microRNAome plays an important role in local mRNA availability. We detected a number of hub microRNAs and especially micoRNA-27b-3p, microRNA-22-3p, the cluster consisting let-7b-5p, let-7c-5p and let-7i-5p as wells as microRNA-181a-5p, microRNA-9-5p and microRNA-124-5p appear as central regulators of the synaptic mRNA pool **(Fig 1F).**

**Figure 1:**
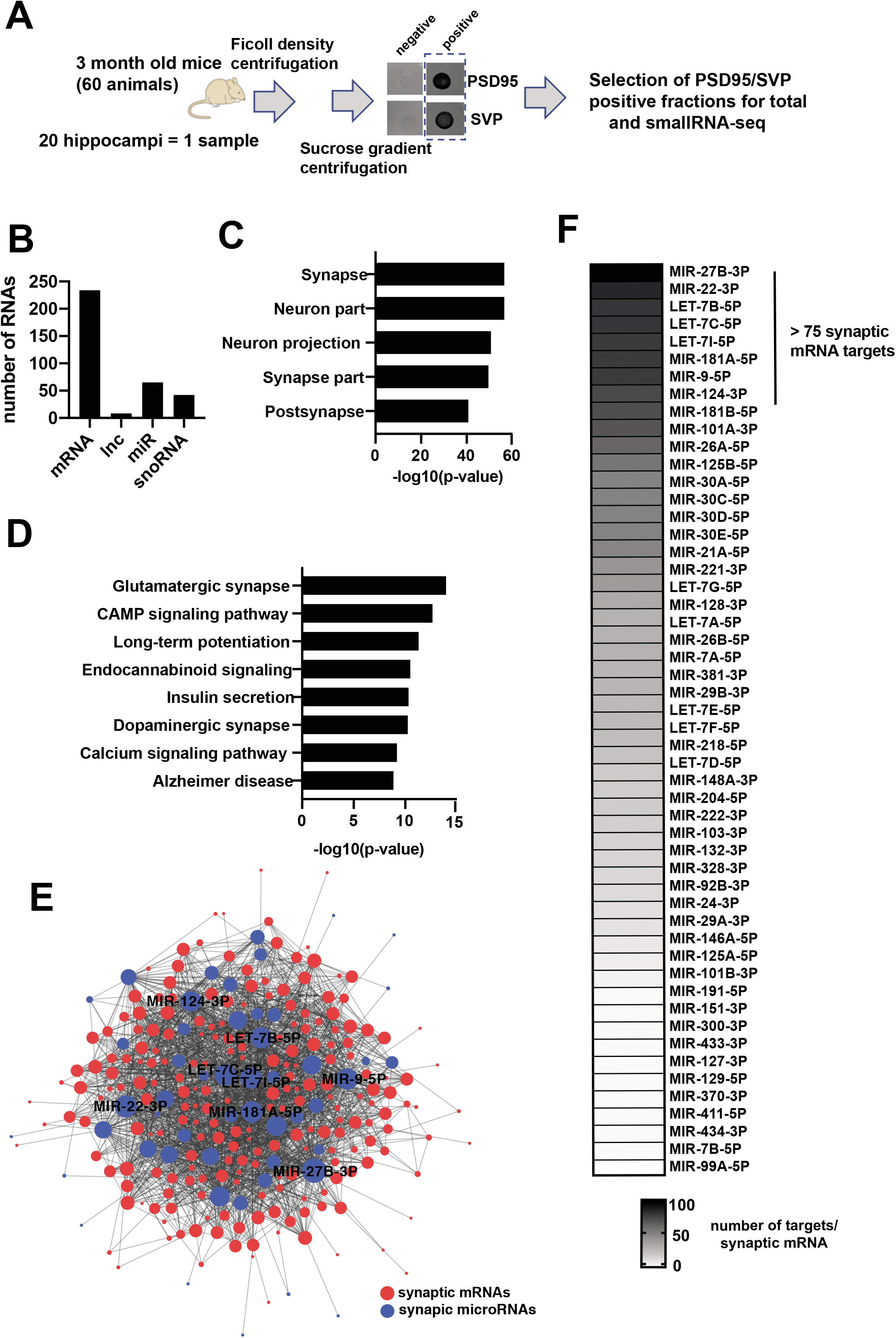
The coding and small-non-coding RNAome of hippocampal synaptosomes. **A.** Experimental scheme. **B.** Bar graph showing the detected RNA species. **C.** GO-analysis showing that the identified mRNAs represent the synaptic compartment. D. KEGG-pathway analysis showing that the synaptic mRNAome consists of transcripts that are essential for the function of hippocampal synapses. E. microRNA-mRNA interaction network of the synaptic RNAome. Red circles represent the identified mRNAs that form a highly connected network, while blue circles indicate the detected microRNAs. Only the names of the top hub microRNAs are shown. F. Heat map showing the synaptic microRNAome ranked by their confirmed mRNA targets that were found at synapses.

### <u>Comparison of the hippocampal synaptosomal</u> coding and non-coding RNAome to primary hippocampal neurons

Compartmentalized microfluidic chambers have been developed to study the pre- and postsynaptic compartments of neurons. In these chambers, neurons grow their neurites into microgrooves and form synapses in a narrow compartment, the perfusion channel (Taylor *et al.*, 2010). We hypothesized that such microfluidic chambers would be a *bona fide* complementary approach to study the synaptic RNAome via RNA-sequencing. Moreover, since the synapses formed within the perfusion channel of such chambers are not in contact with any other neural cell type, this approach would also allow us to address the question to what extent synaptically localized RNAs originate from the corresponding cell or may have been shuttled from neighboring glia cells, a process that has been specifically proposed for synaptic microRNAs (Prada *et al*, 2018). However, a reliable approach to isolate synapses and corresponding RNA for subsequent sequencing from the perfusion channel of such microfluidic chambers did not exist. Therefore, we generated a modified microfluidic chamber <u>that allowed us to</u> cut the perfusion channel to harbor the corresponding synapses followed by the isolation of RNA. Thus, we grew mouse hippocampal neurons in these chambers **(Fig 2A).** For <u>reproducible</u> cutting we employed a newly-devised instrument we call SNIDER (SyNapse Isolation DevicE by Refined Cutting) **(Fig 2B, C)** and isolated RNA for total and smallRNA sequencing. When comparing the transcriptome obtained from the perfusion channel, with corresponding data generated from RNA isolated from primary hippocampal neurons grown in normal culture dishes, we observed the expected enrichment for a specific subset of RNAs, representing about 12% of the entire transcriptome **(Fig 2D).** In more detail, the transcriptome of the perfusion chamber consisted of 1460 mRNAs, 199 lncRNAs, 54 microRNAs and 57 <u>highly expressed snoRNAs of which 22 were also detected in synaptosomes</u>. **(Fig 2E, Supplemental tables S4, 5, 6).** GO-term analysis revealed that the identified mRNAs represent the synaptic compartment, which is in line with our data obtained from the adult mouse hippocampus **(Fig 2F)** and further supports the feasibility of our approach. Functional pathway analysis confirmed that the detected mRNAs code for key synaptic pathways and reflect the high energy demand of synapses (oxidative phosphorylation). This is also the reason why pathways such as Alzheimer’s, Huntington’s and Parkinson’s disease are identified **(Fig 2G),** since key genes de-regulated in these diseases are linked to mitochondria function. The direct comparison of the hippocampal synaptic mRNAome from the adult mouse brain and the mRNAome from primary neurons revealed that almost all mRNAs detected from *in vivo* synaptosomes, are also found in primary neurons grown in microfluidic chambers **(Fig 2H),** confirming 219 mRNAs as a high-quality and reproducible synaptic mRNAome. The GO-terms and functional pathways linked to these 219 mRNAs are identical to the data shown in Fig 1C&D. The 1244 mRNAs that were specifically observed in microfluidic chambers represent also the synaptic compartments and pathways linked to oxidative phosphorylation, synaptic vesicle cycle and metabolic processes and may therefore reflect the difference of the synaptic RNAome in the adult brain and cultured primary neurons **(Fig 2I).** In addition, “neuronal projection” is detected as a significant GO-term, most likely indicating the fact that unlike synaptosomal preparations, the perfusion channel still contains some neurites. <u>This might also explain that much more lncRNA, namely 199 annotated lncRNAs, are detected in the microfluidic chambers. Pathway analysis suggest that these lncRNA are mainly linked to mRNAs that control processes associated with oxidative phosphorylation and synaptic plasticity while comparatively few microRNAs seem to be regulated by the synaptic lncRNAs (**Fig S2)**</u>. Similar to the *in vivo* data, we found 54 highly expressed microRNAs **(Table S5).** To further study the mRNA/microRNA network, we used the same approach as described for the synaptosomal data. Our data reveals that the 88% of the synaptic mRNAome in microfluidic chambers is targeted by 45 (83%) synaptic microRNAs **(Fig 3A).** Taken together, our data from hippocampal synaptosomes and the novel microfluidic chamber strongly suggest that the synaptic transcriptome is under tight control of a local microRNA network. Comparison of the *in vivo* synaptic microRNAome to the data obtained from the microfluidic chambers revealed 17 microRNAs that were commonly identified at synapses, while 37 microRNAs were specific to the chambers and 48 microRNAs were only found in the *in vivo* data from hippocampal synaptosomes **(Fig 3B).** When we generated the synaptic microRNA/mRNA network for the commonly detected 17 synaptic microRNAs and 219 mRNAs (see Fig 2G), we observed that this core synaptic microRNAome controls 80 % (179 of 219) of the core mRNAome **(Fig 3C).**

**Figure 2:**
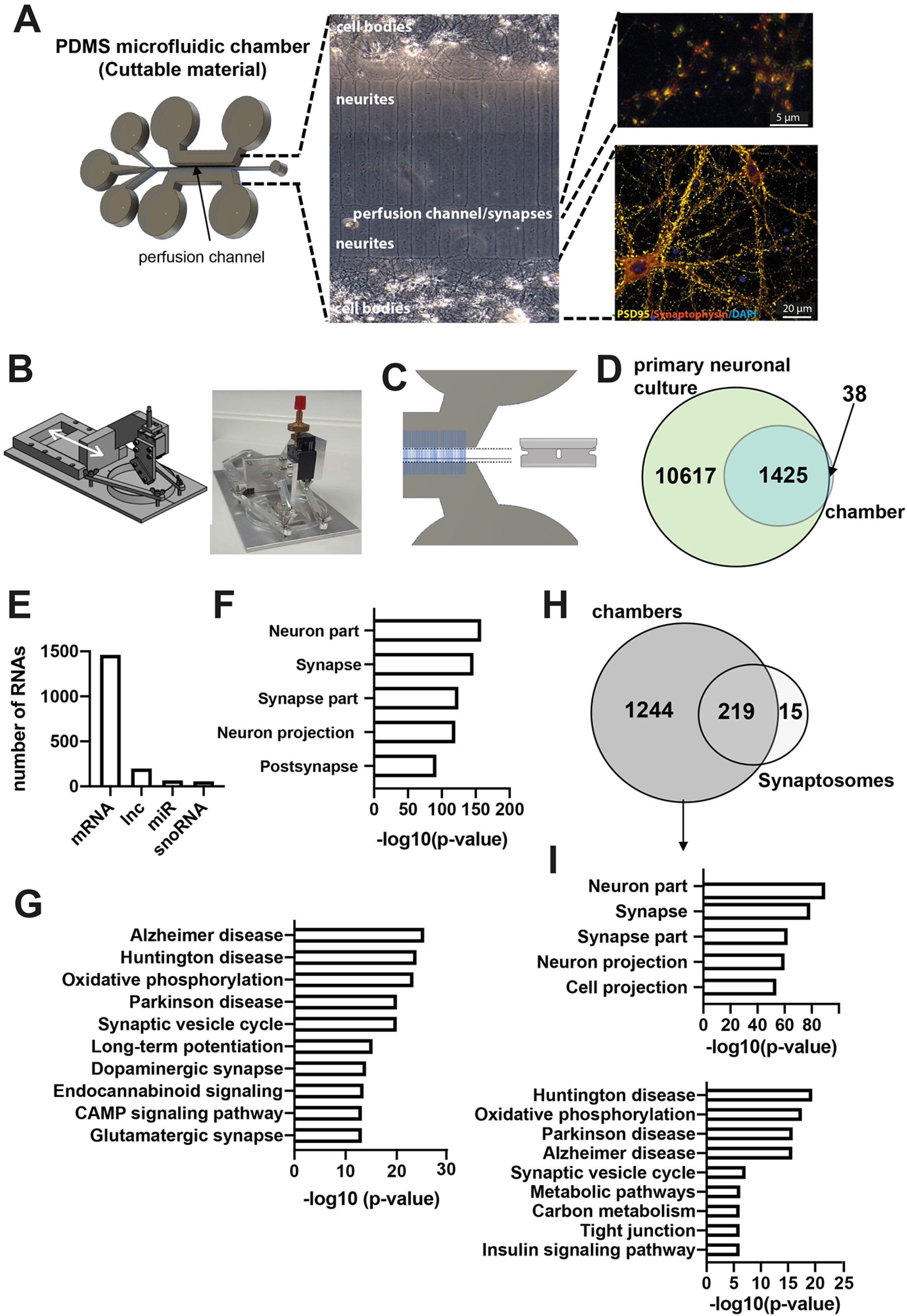
Analyzing the synaptic RNAome in microfluidic chambers via RNA-sequencing. **A.** Microfluid chambers build from PDMS. Left panel shows the scheme of the microfluidic chamber indicating the perfusion channel in which most of synapses form. The principle is based on chambers first reported by Taylor and colleagues (Taylor *et al.*, 2010) but has been substantially modified (See Fig S1 for more details). The middle panel shows the bright-field image of neurons growing in these chambers and the right panel shows immunostaining for PSD-95 and Synaptophysin within the perfusion channel (upper image) and the part of the chambers that contains the cell bodies (lower image). B Scheme and image showing our newly devised tool for cutting the perfusion channel from the microfluidic chambers, named SNIDER. C. Schematic illustration of the cutting of the microfluidic chambers. D. Venn diagram showing the comparison of the total RNA-seq data obtained from primary hippocampal cultures grown in normal dishes (primary neuronal culture) and corresponding data obtained from the perfusion channel isolated from microfluidic chambers in which primary hippocampal neurons were grown. E. Bar chart showing the detected RNA species. F. GO-analysis showing that the identified mRNAs represent the synaptic compartment. **G.** KEGG-pathway analysis showing that the synaptic mRNAome consists of transcripts that are essential for the function of hippocampal synapses. H. Venn diagram showing the overlap of mRNAs detected in hippocampal synaptosomes and in microfluidic chambers. **I.** Upper panel: GO-analysis showing that the 1244 mRNAs specifically detected in microfluidic chambers represent the synaptic compartment and “cell projection”. Lower panel: KEGG-pathway analysis of the same dataset.

**Figure 3:**
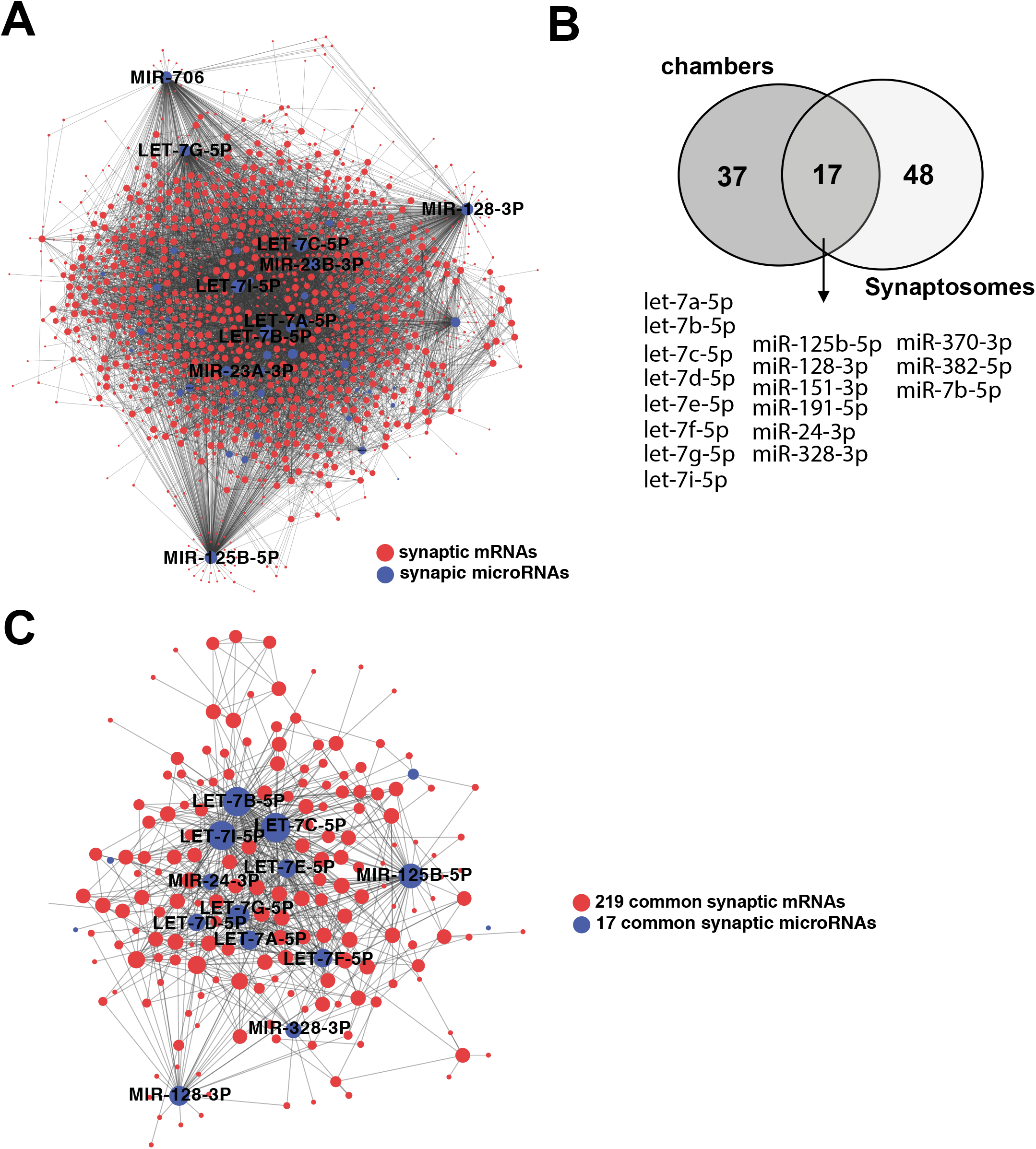
A core synaptic microRNAome. **A.** microRNA-mRNA interaction network of the synaptic RNAome detected in microfluidic chambers. Red circles represent the identified mRNAs that form a highly connected network, while blue circles indicate the detected microRNAs that control this network. Only the names of the top hub microRNAs are shown. B. Venn diagram comparing microRNAs detected in microfluidic chambers (Chambers) and synaptosomes. C. microRNA-mRNA interaction network of the 219 synaptic mRNAs commonly detected in synaptosomes and microfluidic chambers and the 17 commonly detected microRNAs. Only the names of the top hub microRNAs are shown.

### Evidence for astrocytic microRNA transport to synapses

The finding that 37 microRNAs are exclusively found in synapses from primary neurons is likely due to the difference between *in vivo* brain tissue and primary neuronal cultures and a similar trend has been observed at the level of the mRNAs **(see Fig 2G).** More interesting is the observation that 73%, namely 48 out 65, of the microRNAs detected in hippocampal synaptosomes are not found in microfluidic chambers (see Fig 3B), which is in contrast to the mRNA data in which almost all of the synaptosomal mRNAs are also found in synapses of the primary neuronal cultures <u>grown</u> in microfluidic chambers (See Fig 2G). These data may indicate that *in vivo* some of the synaptic microRNAs are not exclusively produced by the corresponding neuron but might be rather shuttled to synapses via other neural cell types. In fact, movement of microRNAs between cells is an accepted mechanism of intra-cellular communication (Jose, 2015). Prime candidate cells to support synapses with microRNAs are astrocytes that form together with neurons tripartite synapses. A prominent mechanism that mediates RNA transport amongst neuronal cells is intracellular transport via exosomes (Smythies & Edelstein, 2013). Thus, we compared a previously published dataset in which microRNAs from astrocytic exosomes were analyzed via a TAQman microRNA-array (Jovičić *et al*, 2013). Indeed, 50% of the microRNAs exclusively detected in hippocampal synaptosomes have also been described in exosomes released from astrocytes **(Fig 4A).** When we asked if these 23 microRNAs have mRNA targets detected in synaptosomes we observed that 21 of these microRNAs target in total 197 out of the commonly detected 219 synaptic RNAs **(Fig. 4B),** which is further confirmed by functional pathway analysis showing that the 21 microRNAs control synaptic genes linked to the glutaminergic synapse, LTP and cAMP signaling **(Fig. 4C).** It is interesting to note that the synaptic mRNAs not targeted by any of the 21 microRNAs represented functional pathways linked to oxidative phosphorylation **(Fig. 4D).**

**Figure 4:**
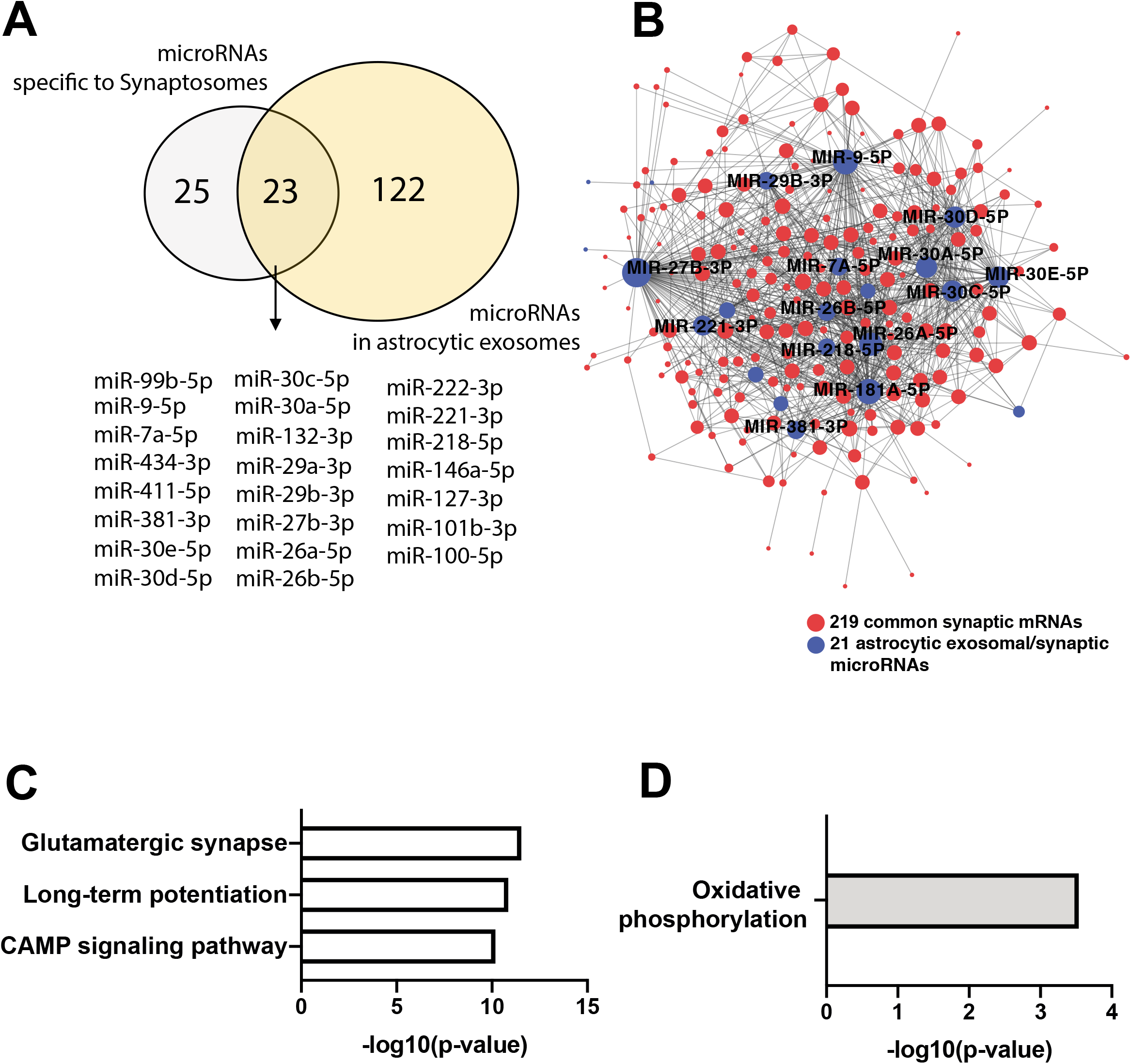
Comparing microRNAs from astrocytic exosomes to the synaptic RNAome. **A.** Venn diagram comparing the 48 microRNAs exclusively detected in synaptosomes to the list of microRNAs found in astrocytic exosomes. B. microRNA-mRNA interaction network showing that 203 of the commonly detected 219 mRNAs and 21 of the 23 microRNAs found in synaptosomes and astrocytic exosomes form an interaction network. Only the names of the top hub microRNAs are shown. C. KEGG-pathway analysis of the 203 mRNAs within the network. D. KEGG pathway analysis of the 16 common synaptic mRNAs that are not targeted by the overlapping microRNAs shown in (A).

### Synaptic microRNAs are linked to neurodegenerative and neuropsychiatric diseases

So far, our data support the view that the synaptic microRNAome plays an important role in neuronal function. To further strengthen this notion, we decided to ask whether synaptic microRNAs might be particularly de-regulated in cognitive diseases. To this end we performed a literature search and curated a list of 71 microRNAs that were found to be de-regulated in post-mortem human brain tissue, blood samples or model systems for Alzheimer’s disease, depression, bi-polar disease or schizophrenia. Comparison of this dataset with our findings from synaptosomes revealed 17 synaptic microRNAs that are de-regulated during cognitive diseases of which 4 are also found in the microfluidic chambers and 11 were also detected in astrocytic exosomes, representing an interesting pool of synaptic microRNAs for further studies **(Table 1).**

### Microfluidic chambers are suitable to assay the synaptic RNAome upon neuronal stimulation

Our findings suggest that we can study the synaptic RNAome in a reliable manner using our modified microfluid chambers in combination with SNIDER. This approach also provides a novel tool to study the neuronal-controlled synaptic RNAome in response to stimulation. To further evaluate this potential, we decided to expose primary hippocampal neurons grown in microfluidic chambers to KCl treatment followed by the isolation of the perfusion channel and RNA isolation for RNA-sequencing 2 h later **(Fig 5A).** Our analysis revealed a substantial number of mRNAs that were increased in the synaptic compartment **(Fig 5B, Table S7).** Since we can exclude that these mRNAs are shuttled from glia cells, they likely represent part of the transcriptional response and reflect mRNAs that were transported to synapses, which is feasible within the 2h time window after treatment. In line with this assumption the up-regulated RNAs exclusively represent the synaptic compartment **(Fig 5C).** Functional pathway analysis revealed a strong enrichment of RNAs coding for the ribosome **(Fig 5D).** In fact, 50% of all transcripts that correspond to the ribosomal subunits were increased at the synapse upon KCL treatment **(Fig 5E).**

**Figure 5:**
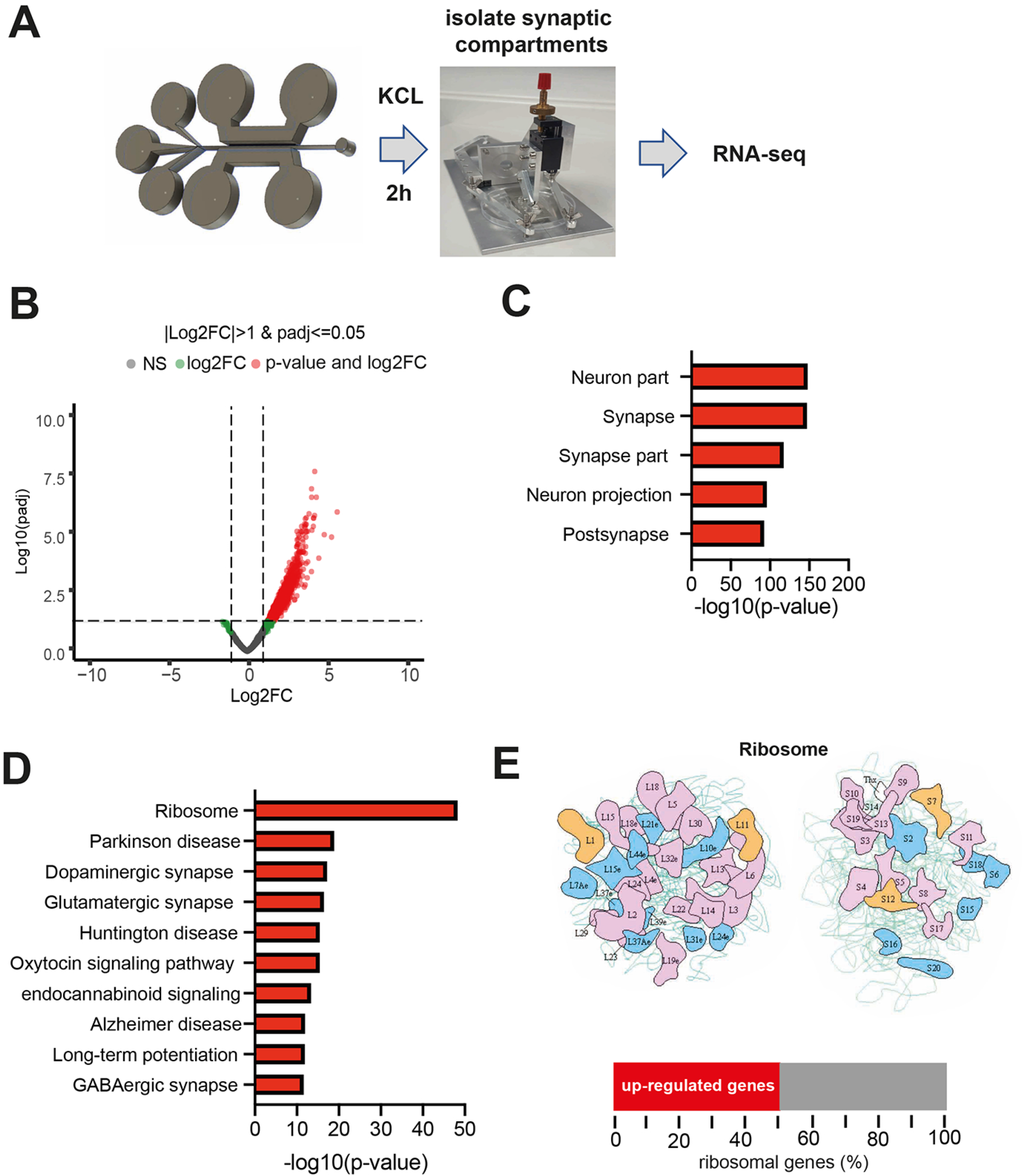
The synaptic mRNAome upon stimulation. **A.** Experimental scheme. **B.** Volcano plot showing a substantial up-regulation of synaptic RNAs upon KCL treatment. **C.** GO-analysis showing that the identified mRNAs represent the synaptic compartment. D. KEGG-pathway analysis showing that the changes of the synaptic mRNAome upon KCL treatment represent transcripts mainly linked to ribosomal function. E. Upper panel shows images of the KEGG pathway for “ribosome”. Colored subunits represent transcripts significantly increased. Lower panel: bar chart showing that 50% of the genes that comprise the “ribosome” KEGG-pathway are increased at the synaptic compartment upon KCL treatment.

## Discussion

### The synaptic RNAome

The aim of our study was to provide a high-quality dataset of the synaptic coding and small non-coding RNAome with a specific focus on microRNAs. Thus, our data represents an important resource for future studies. To the best of our knowledge our study also provides the first dataset which analyzes in parallel the coding, <u>non-coding</u> and small non-coding RNAome in hippocampal synapses via next-generation sequencing. Moreover, we used two different approaches in that we isolate hippocampal synaptosomes from the hippocampus of 3 months-old wild type mice and we developed a microfluidic chamber that in combination with a novel cutting device allowed us to isolate synaptic compartments for subsequent RNA-sequencing from primary hippocampal neurons. This chamber combines the advantages of the currently used microfluidic chambers that allow the specific manipulation of synapses (Taylor *et al.*, 2010), with the ability to isolate the perfusion channel that harbors synaptic connections. Therefore, this novel microfluidic chamber will allow the specific manipulation of the synaptic compartment in combination with next-generation sequencing approaches and should be viewed as a suitable screening tool to study the dynamics of the synaptic RNAome. Feasibility of this approach was for example demonstrated by our finding that KCL treatment leads to substantial changes of the synaptic RNAome and future approaches will now employ more physiological manipulations and study the synaptic RNAome in disease models. It is noteworthy, that most of the mRNAs up-regulated at the synapse upon stimulation represent key components of the ribosome, which is in agreement with the importance of local mRNA translation (Holt *et al.*, 2019).

In line with previous data we identified a substantial number of mRNAs that almost exclusively represent the synaptic compartment and key signaling pathways linked to synaptic integrity and plasticity. <u>It is interesting to note that the mRNA coding for the amyloid-precursor protein (APP), a key factor in Alzheimer’s disease (AD) pathogenesis, was also found at synapses **(see Table S1).** To our knowledge, this observation has not been explicitly reported before but is in line with the physiological function of wild-type APP at synapses (Hefter *et al*, 2020)</u>. Generally, we detected more mRNAs within the dataset obtained from primary neurons when compared to the synaptosomal preparation. This observation likely reflects the difference between the *in vivo* preparation of hippocampal tissue and cultured primary neurons. Another important consideration is that synapses likely differ depending on the distance to the soma, an issue that cannot be addressed when isolating synaptosomes, while the RNAome detected in the microfluidic chambers represent synapses that are most distant to the corresponding somata. Similar important is the fact that the preparation from the perfusion channel of our microfluid chamber still contains some neurites. Thus, the corresponding RNAome also includes dendritic mRNAs. This view is supported by previous data in which the mRNA pool was analyzed from neuropil or synapto-dendritic compartments. For example, 2550 mRNAs were detected in hippocampal neuropil from mice (Cajigas *et al.*, 2012), and 1875 mRNAs where identified when ribosome-bound mRNA was analyzed in the same region (Ainsley *et al.*, 2014). We observed only 234 mRNAs in hippocampal synaptosomes but we suggest that these mRNAs represent a high-quality dataset. Thus, we only report mRNAs that passed rigorous quality control and exhibit a substantial amount of sequencing reads. The quality of these data is further confirmed by the fact that almost all of the synaptosomal mRNAs, namely 219, are also detected within the RNA-seq dataset we obtained from the microfluidic chambers. The most comparable mRNA dataset to our *in vivo* approach is a recent study that employed FACS to isolate synaptosomes from the mouse forebrain (Hafner, 2019) and also reported raw data on the generic synaptosomes. It is important to note that this study employed a different analysis pipeline and reported all transcripts that map with >25% of the read length when using the STAR-aligner tool, while we consider only transcripts that map with at <u>least >66%</u>. Nevertheless, we observed that the top 500 mRNAs reported by Hafner et al. almost completely overlapped with our dataset. Namely 209 of the 234 mRNAs that we reported for hippocampal synaptosomes are also found in the Hafner et al. dataset from mouse forebrain synaptosomes **(table S8),** further supporting the quality of our dataset and strengthening the view that synaptic mRNAs play a critical role in neuronal function. We also report the detection of lncRNAs in datasets obtained from synaptosomes and microfluidic chambers but for now restricted the presented data to the currently annotated lncRNAs. We also detected <u>lncRNAs that are currently still referred to as “predicted” and await further confirmation. Therefore, we encourage researchers to further explore our raw data as annotation of the genome improves</u>. The presence of lncRNA in synaptosomes is in line with previous data (Chen *et al.*, 2017) but it is interesting to note that more lncRNAs were found in the microfluidic chambers when compared to the *in vivo* synaptosomes. <u>A similar trend has been observed for mRNAs and might be due to the fact that the RNA preparation from the microfluid chambers also contain some dendritic RNA. Our data suggest that the detected lncRNAs regulate processes associated with oxidative phosphorylation and synaptic plasticity and may also affected the function of selected microRNAs. Although these observations need to be further studied, it is interesting to note that metastasis-associated lung adenocarcinoma transcript 1 (Malat1) appeared as one hub lncRNA at synapses. This is in line with a previous study showing that knocking down MALAT1 in hippocampal neurons decreases the number of synapses, although it has to be mentioned that the authors linked this finding to the role of MALAT1 on gene-expression control (Bernard *et al*, 2010)</u>. The presence of snoRNAs at synapses is also highly interesting and in line with a previous study that reported snoRNAs in synaptosomes (Smalheiser *et al*, 2014). Moreover, there was a substantial overlap of the snoRNAs detected in synaptosomes and in primary <u>neurons (60% of the synaptosomal snoRNAs were also detected in microfluidic chambers). Most of the commonly detected snoRNAs</u> were of the C/D box <u>(49%)</u> or H/ACA-box type <u>(17%)</u> that regulate RNA-methylation and pseudouridylation of mainly ribosomal RNAs (Bratkovič *et al*, 2020), which is in line with the presence of ribosomes at synapses (Holt *et al.*, 2019). <u>However, we also identified snoRNAs that cannot be classified in either category (35%) that warrant further investigation</u>. Some of the synaptic snoRNAs have been associated to additional processes and for example SNORD50, SNORD83B or SNOR27 have been linked to mRNA 3’ processing and post-transcriptionally gene-silencing (Bratkovič *et al.*, 2020), while SNORD115 affects mRNA abundance and is genetically linked to the Prader-Willi-syndrome, a rare genetic disease leading to intellectual disability (Cavaillé, 2017).

### A synaptic mRNA/microRNA network

We detected a substantial number of microRNAs in hippocampal synaptosomes and in the microfluidic chambers. The presence of mature microRNAs at synapses is in line with previous reports that employed RT-PCR to study neurites of primary hippocampal neurons (Kye *et al*, 2007), micro-array-technology to analyze microRNAs in the synapto-neurosomes isolated from the forebrain of mice (Lugli *et al*, 2008) or more recently also smallRNA-sequencing and NanoString analysis of hippocampal neuropil or synaptosomes (Smalheiser *et al.*, 2014) (Sambandan *et al.*, 2017). Comparison of the dataset generated by Sambandan and colleagues revealed that out of the 65 microRNAs we detect, 57 were also reported in this previous study. These data further strengthen the view that microRNAs play an important role at synapses and suggest that our dataset represents a high quality synaptic microRNAome as a resource for future studies. To the best of our knowledge, our study is the first that provides a synaptic coding and small non-coding RNAome from the same preparation thereby allowing us the address the role of the synaptic microRNAome at the systems level. We used the data to develop a novel tool which is first fed with the mRNA data to parse multiple databases containing experimentally validated interactions and thereby building a high confidence mRNA network of the synapse (See methods for more details). We intersected this mRNA network with the confirmed targets of all microRNAs, which are detected within the same sample to build the synaptic microRNA/mRNA network. Overall, our data suggest that up to 98% of the synaptic mRNAome is controlled by synaptic microRNAs, suggesting that essentially all synaptic localized mRNAs are potentially regulated via the synaptic microRNAs. Considering that mRNA transport to synapses is an energy-demanding and highly controlled process (Doyle & Kiebler, 2011) it is likely that synaptic microRNAs do not degrade their mRNA targets but rather control their availability for local translation, a question that should be studied in future experiments at the systems level. Another important observation is that many of the synaptic microRNAs are de-regulated in cognitive diseases (see table 1) that often start with synaptic dysfunction. In addition, there is increasing interest in circulating microRNAs as biomarkers for cognitive diseases (Rao *et al*, 2013) (Rupaimoole & Slack, 2017). The fact that microRNAs have also been reported in synaptic vesicles (Xu *et al*, 2013) and in exosomes derived from neuronal cultures (Jain *et al*, 2019) suggest a potential path how pathological microRNA changes observed in the brain may also manifest in circulation. Hence, the various CNS clearance systems (Plog & Nedergaard, 2018) might transport such vesicles to the circulation, a hypothesis that should be further studied. In the same context, there is substantial data to suggest that microRNAs regulate biological processes across cell-types and even organs (Valadi H, 2007) (Jose, 2015). Intriguingly, in the perfusion channel of microfluidic chambers, which are free of any somata and only contain distal synapses and some neurites, substantially less microRNAs are existent than in the synaptosomes. These microRNAs significantly overlapped with the ones detected in exosomes released by astrocytes (Jovičić *et al.*, 2013). It is therefore tempting to speculate that within the tripartite synapse astrocytes support synapses with additional microRNAs that help to control the synaptic mRNA pool. Support for this view stems also from the observation that the 3 most significant functional pathways controlled by the synaptosomal microRNAome are “Glutamatergic synapse”, “cAMP signaling” and “long-term potentiation”, which are identical to the top 3 pathways controlled by the microRNAs that are potentially shuttled to synapses via astrocytes. These data underscore the importance of the corresponding mRNA pool and may suggest that microRNAs supplied to synapses by other cell types might suppress translation of the most relevant local mRNAs rather than degrading a few selected RNAs. Our data allowed us to identify a number of synaptic hub microRNAs (e.g. see Fig 1E and F) and the functional analysis of these microRNAs would be an important task for future studies. Of particular importance would be microRNAs that are de-regulated in cognitive diseases. Support for this view stems from recent data on microRNA-181a-5p, a hub in our synaptic network, that is de-regulated in neurodegenerative and neuropsychiatric diseases (Stepniak *et al*, 2015) (Ansari, 2019) and was found to be processed at synapses upon neuronal activity (Sambandan *et al*, 2017). The finding that most microRNAs of the let-7 family are highly abundant at synapses and control a large set of mRNAs is also interesting is interesting, since these microRNAs have been observed in several CNS-related pathologies (Derkow *et al*, 2018) while comparatively little is known on their role on the adult brain. Another hub microRNAs is miR-125b-5p that is de-regulated in Alzheimer’s disease and causes memory impairment in mice when elevated in the hippocampus of mice (Banzhaf-Strathmann *et al*, 2014), yet its role at the synapse remains elusive. Similarly interesting is miR-128-3p, that is de-regulated in various neuropsychiatric and neurodegenerative diseases and recent data suggest that inhibition of microRNA-128-3p can ameliorate AD pathology (Liu *et al*, 2019).

In conclusion, our study provides the synaptic RNAome and is thus a valuable resource for future studies. Our data furthermore support the importance of synaptic mRNAs and microRNAs and we introduce a new microfluidic chamber that will allow researchers to combine the power of a specific analysis and manipulation of the synaptic compartment (Taylor *et al.*, 2010) with RNA-sequencing approaches.

## Materials & Methods

### Animals

Three months old male C57B/6J mice were purchased from Janvier Labs. All animals were housed in standard cages on 12h/12h light/dark cycle with food and water ad libitum. All experiments were performed according to the protocols approved by local ethics committee

### Isolation of hippocampal synaptosomes for RNA-sequencing

<u>To obtain sufficient RNA for sequencing of hippocampal synaptosomes we isolated the hippocampi from sixty 3-month-old wild type mice. Twenty bi-lateral hippocampi were pooled as 1 sample to obtain 3 independent samples that were further processed to isolate high-quality synaptosomes u</u>sing a previously described protocol (Boyken *et al*, 2013)._ln brief, hippocampi were homogenized by 9 strokes at 900 rpm in sucrose buffer and centrifuged at 4° for 2min at 5000rpm (SS34). Supernatants were further centrifuged at 4° for 12min at 11000rpm. Pellets were loaded onto a Ficoll gradient and centrifuged at 4° for 35min at 22500rpm (SW41). The interface between 13% and 9% Ficoll was washed by further centrifugation and then pelleted by 8700rpm for 12min in a SS34 rotor. Resuspended synaptosomes were then centrifuged on a sucrose gradient for 3h at 28000rpm (SW28). Finally, synaptosomes were fractioned via the Gilson Minipuls and 21 fractions were collected and analyzed by dot blotting. For this, from each fraction, 2μl of sample were pipetted on nitrocellulose membrane, and dried for 5min. Blocking of unspecific signal was done by 5% low fat milk in TBST for 10 min. Antibodies against Synaptophysin and PSD95 were applied for 15min, then the membrane was washed three times for 3min each, in TBST with 5% milk. Secondary antibody was applied for 15min. Afterwards membrane was washed again three times with TBST without milk before being imaged. <u>Only 5 fractions from each preparation showed a signal for synaptophysin and PSD95 ensuring the presence of high-quality synaptosomes and were therefore processed for total and small RNA-sequencing</u>.

### Production of microfluidic chambers

<u>To isolate synapses and corresponding RNA for subsequent sequencing from the perfusion channel of currently employed microfluidic chambers (Taylor *et al.*, 2010) was not possible. Therefore, we generated a microfluidic chamber that allowed us to cut the perfusion channel by using polvdemethvlsiloxan (PDMS) for the chamber and the corresponding substrate **(Fig S1).** Pilot studies showed that unlike the commonly used microfluidic chambers (Taylor *et al.*, 2010), the usage of PDMS as a substrate to bind the chambers on allowed us to cut the perfusion channel. In more detail</u>, the microfluidic chambers were designed using AutoCAD 2017. The overall layout was similar to the version reported by Taylor and colleagues (Taylor *et al*.), yet for more yield of synaptic RNAs the length of the chamber was increased, with more microgrooves and a wider synaptic compartment to allow easier alignment during cutting. Layouts were translated into photolithography masks by Selba. Production of silicon wafers was done with two layers. The first layer was made by applying 2 ml Photoresist SU-8-2025 on 50.8mm diameter silicon wafers and running the spin coater with the following settings: 1.) 15 sec, 500 rpm, 100 ramp 2.) 100sec, 4000 rpm, 50 ramp. To prebake, wafers were put on a 65° heating plate for 1 min, then for 15min on a 95° heating plate. For depositing the first layer, the mask with the microgrooves pattern was inserted into the MJB4 mask aligner; exposure was set to 9 sec under light vacuum conditions. Afterwards wafers were postbaked at 65° for 1 min and 5 min at 95°C.

Subsequently 3 ml of the second photoresist SU-8-2050 were added on top and spread thin with the following spincoater protocol: 1.) 15 sec, 100 ramp, 500 rpm 2) 60 sec, 900 rpm, 50ramp. This time prebaking was done with 1min at 65° and minimum of 30min at 95°. The second layer was aligned to the microgrooves using the microscope of the mask aligner. UV light exposure lasted 19 sec, in the soft contact setting. After postbaking as described for the first layer, wafers were developed for 10min or more in mrDev600 with the aid of ultrasonication. PDMS (SYLGARD™ 184 Silicone Elastomer Kit) was used to manufacture the chambers as well as the bottom substrates. Sylgard components were mixed 10:1, mixed with a 1ml pipette tip, poured over the wafers that were placed in 6cm diameter Petri dishes and very thinly (1-2mm high) onto 10cm dishes. Degassing was done for minimum 15 minutes in a desiccator under vacuum. Afterwards wafers and bottom parts were transferred to a 70° oven and cured for 2h. Chambers and bottom parts were cut out by a scalpel, holes in the chambers were punched by biopsy punchers of 6mm and 8mm diameter and bottom parts were cut into smaller pieces to hold one chamber each. To clean off dust, the pieces off of dust they were placed in an ultrasonic bath for 10 min and then dried on a heatplate at 70°. PDMS can be bound to PDMS covalently under oxygen plasma conditions; a tesla-coil type device, the Corona plasma treater from Blackhole lab, was used to this end. The plasma treater was hovered slowly 2cm above the chambers (bottom side up), going back and forth to cover the whole area by discharges for 30sec, then the same was done to the bottom part. Thereupon both parts were brought together and pressed very slightly to ensure complete contact. Covalent bond forming was enhanced by placing the so assembled chamber in the oven at 70° for 10min. Subsequently chambers were filled with PBS or borate buffer to maintain hydrophilic properties. For chambers that were supposed to be imaged, chambers were not treated with plasma; rather chambers were assembled to the PDMS or glass substrate under the biosafety cabinet by simply pressing both pieces together. Once assembled, chambers were brought to a biosafety cabinet and washed with 70% ethanol, then twice with water. Coating on PDMS worked best when done with 0.5 mg PDL in borate buffer overnight. Visual inspection under the microscope should make sure that no bubbles are present in the chambers. Great care needs to be taken when washing to not remove the coating. Liquid should be never removed with a suction pump sucking liquid directly from the channels, instead liquid should be removed by pointing the pipette at the wall of open reservoirs. Washing was done twice with PBS, 80μl per top reservoir, allowing for the liquid to flow into the down reservoir. Perfusion reservoirs were washed by applying 50μl in each well, one at a time and waiting for 5min in between. Once all PBS was removed from the open reservoirs 80μl of medium was added per top reservoir, allowing for the liquid to flow into the down reservoir. This process was repeated once, before chambers were left over night in the incubator before seeding, to ensure proper hydrophilicity. For easier handling always two chambers were put together in a 10cm dish, with two lids of 15ml falcon tubes filled with water next to them, to reduce the evaporation from the chambers themselves.

### Primary hippocampal neuronal cultures

Pregnant CD1 mice were sacrificed under anesthesia by cervical dislocation at E16 or E17. Brains from embryos were extracted and their hippocampi collected. Processing was done using the Papain kit from Worthington, and cells were counted and diluted to a density of 5 Million per ml. Seeding was done with the following pipetting scheme in order to make sure, most cells reach the microgrooves but do not enter them, 10μl of cell suspension containing 70.000 cells were injected in the channels from the top wells. We started with the axonal side. A second pipetting step with 5μl added to the channels from the bottom wells, after inspection of cells under the microscope. After 10 minutes, a similar seeding was performed for the dendritic side. One hour later each well was filled up to 100μl. The next day another 100μl were pipetted into each well. Visual inspection under a microscope was necessary to do several rounds of seeding with decreasing volume to made sure the desired spread of cells was achieved. After two hours reservoirs of the chambers were filled up with medium to 100μl each, by pipetting an additional 70μl simultaneously in both reservoirs per side, while not adding more medium to the perfusion. We used Neurobasal Plus with GlutaMax, Penicilin/Strep and B27 Plus supplement for better viability. Parallel to chambers, normal 12-well dishes, coated with PDL in borate overnight and washed three times with water, were cultured at 260.000 cells per well; those served as standby cultures. Since medium evaporation can happen quickly in the chambers, every 2-3 days medium from these standby cultures was filtered by a 0.22μm syringe filter and then added to the chambers. For the KCl stimulation, around 50μl of medium was collected from each reservoir of the chambers, mixed with KCl as to result in a final concentration of 50mM when given back to the chamber and then incubated for 2h before RNA isolation.

### Harvesting of synaptic RNAs from microfluidic chambers: SNIDER

In order to parallel cut the PDMS substrate, we designed a machine consisting of a blade-holding arm on a ball-bearing rail, allowing frictionless mobility in one dimension. A screw-driven spring drives the razorblades height position and allows for controlling the penetration depth of the blades into the PDMS. The non-cutting corners of the razorblade were removed with a plunger to only have one accessing point of the blades into the PDMS. Small metal plates were put in between the blades and served as spacers, increasing the inter-blade distance to 900 μm. On the day of harvest cells in the chambers were washed once with PDMS and flipped upside down. Great care was taken to maintain a RNAse free environment by prior cleaning of all tools and instruments with RNAsezap and 70% EtOH afterwards. To have an endpoint for the long parallel cut we introduced with a scalpel two horizontal cuts between the outer perfusion wells and the upper left respectively upper right well that met at the perfusion stream. Then chambers were aligned by their perfusion stream on a marked line of the device. By close visual inspection the blades were lowered just before entering the PDMS material and blades were brought in parallel to the synaptic compartment. Blades were then lowered 2mm deep into the substrate just before the perfusion outlet and then the metal lever was pulled backwards, moving the blades towards the perfusion wells until the parallel cut met the V-shaped cut induced by scalpel earlier. With a pair of tweezers, the synaptic compartment was taken out and put into cell lysis buffer solution of GenElute Sigma kit, whereupon we followed the manufactures protocol under 1C to isolate total RNA, including small RNAs.

### RNA sequencing

The synaptosomal RNA samples were split into halves; one was further processed to obtain total RNA libraries using the Illumina Truseq total RNA kit, the other half was used for small RNA sequencing using the NEBNext Small RNA Library Prep Kit as described before (Benito *et al*, 2015). For total RNA sequencing of RNA from microfluid chambers, we always pooled two samples and libraries were created with Takara’s SMARTer Stranded Total RNA-Seq Kit v2 - Pico Input Mammalian, small RNA libraries were generated using Takara’s SMARTer smRNA-Seq Kit for Illumina. To verify the library and sequencing procedure we added spike-in RNAs from the QIAseq miRNA Library QC kit prior to library creation.

### Bioinformatic analysis

Sequencing data was processed using a customized in-house software pipeline. Illumina’s conversion software bcl2fastq (v2.20.2) was used for adapter trimming and converting the base calls in the per-cycle BCL files to the per-read FASTQ format from raw images. Quality control of raw sequencing data was performed by using FastQC (v0.11.5). Trimming of 3’ adapters for smallRNASeq data was done using cutadapt (v1.11.0) (https://doi.org/10.14806/ei.17.1.200). The mouse genome version mm10 was used for alignment and annotation of coding and non-coding genes. Small RNAs were annotated using miRBase (Griffiths-Jones, 2006) for miRNAs and snOPY (Yoshihama *et al*, 2013) for snoRNAs. For totalRNASeq reads were aligned using the STAR aligner (v2.5.2b) (Dobin *et al*, 2013)and read counts were generated using featureCounts (v1.5.1) (Liao *et al*, 2014). For smallRNASeq reads were aligned using the mapper.pl script from mirdeep2 (v2.0.1.2) (Friedländer *et al*, 2012) which uses bowtie (v1.1.2) (Langmead & Salzberg, 2012) and read counts were generated with the quantifier.pl script from mirdeep2. All read counts were normalized according to library size to transcript per million (TPM). We used a TPM cutoff of 1000 reads for smallRNAs to make sure that these smallRNAs were considerably detected up to an average raw count of 10 reads. To account for differences in sequencing depth between synaptosomal mRNAs (average of 6mio unique reads per lane) and mRNAs from microfluidic chambers (average of 20mio unique reads per lane) we applied a cutoff of 50 and 100 normalized reads, respectively. Differential expression analysis was performed with the DESeq2 (v1.26.0) R (v3.6.3) package (Love *et al*, 2014), here unwanted variance was removed using RUVSeq (v1.20.0) (Risso *et al*, 2014). Networks were build using Cytoscape (v3.7.2) (Shannon *et al*, 2003) based on automatically created lists of pairwise interactors. We used in-house Python scripts to detect interactions between expressed non-coding RNAs (miRNAs, lncRNAs, or snoRNAs) and coding genes; interaction information was collected from six different databases: NPInter (Teng *et al*, 2020), RegNetwork (Liu *et al*, 2015), Rise (El Fatimy *et al*, 2018), STRING (Szklarczyk *et al*, 2019), TarBase (Karagkouni *et al*, 2018), and TransmiR (Tong *et al*, 2019). All interactions classified as weak (if available) were excluded. The lists of pairwise interactors were loaded into Cytoscape and all nodes connected by only one edge were removed to build the final network, respectively.

### Imaging

Cells were fixed in 4% PFA in PBS plus 1μM MgCl2, 0.1 μM CaCl2 and 120mM Sucrose. Our imaging setup consists of a Leica DMi8 microscope that is equipped with a STEDYcon. Phase contrast images were obtained using the Leica in its normal mode, with the Leica DMi8 software. All other fluorescent images were taken with the STEDYcon in either confocal or STED mode. Antibodies: PSD95 (Merck - MABN 68) and Synaptophysin 1 (Synaptic Systems 101 004), both diluted to 1:400. Secondary antibodies were StarRED (Abberior, STRED-1001-500UG) and Alexa Fluor 633 Anti-Guinea Pig (Invitrogen, A21105) both diluted to 1:400. DAPI was applied for 1min for counterstaining.

## Availability of data

All sequencing data are available via GEO database. <u>GSE159248</u>: https://www.ncbi.nlm.nih.gov/geo/query/acc.cgi?acc=GSE159248

## Code availability

Not applicable

## Compliance with Ethical Standards

The authors declare no conflict of interest. This work includes experiments with mice. All described experiments approved by the local animal care committee.

## Consent to participate

<u>Not applicable since this study does not involve research on human subjects.</u>

## Consent for publication

<u>Not applicable since this study does not involve research on human subjects.</u>

## Funding

This work was supported by the following third-party funds to AF: the ERC consolidator grant DEPICODE (648898), funds from the SFB1286 (B06), the DFG under Germany’s Excellence Strategy - EXC 2067/1 390729940 the and funds from the German Center for Neurodegenerative Diseases (DZNE).

## Author contribution

<u>All authors contributed to the study conception and design</u>. RE conducted cell culture and RNA-sequencing experiments, build microfluidic chambers and analyzed data. DMK performed bioinformatic analysis, TB isolated synaptosomes, GB and SK helped with the generation of microfluidic chambers, RI performed total RNA-sequencing from hippocampal neuronal cultures in normal culture dishes and curated the disease-related list of microRNAs, AF supervised the project and wrote the manuscript.

## Acknowledgments

RE is a members of the international Max Planck Research School (IMPRS) for Neuroscience, Göttingen. We thank Mike Zippert and Reinhard Hildebrandt for the construction of SNIDER. This work was supported by the following third-party funds to AF: the ERC consolidator grant DEPICODE (648898), funds from the German research foundation (DFG) SFB1286 (B06) and funds from the German Center for Neurodegenerative Diseases (DZNE). SK is supported by the DFG via SFB1286 (B02). AF and SK are supported by funds from the DFG under Germany’s Excellence Strategy - EXC 2067/1 390729940.

## Notes

https://www.ncbi.nlm.nih.gov/geo/query/acc.cgi?acc=GSE159248

### Competing Interest Statement

The authors have declared no competing interest.

