## Supplemental Figures 1&2 for "The Coding And Small-Non-Coding Hippocampal Synaptic RNAome"

**Figure S1**

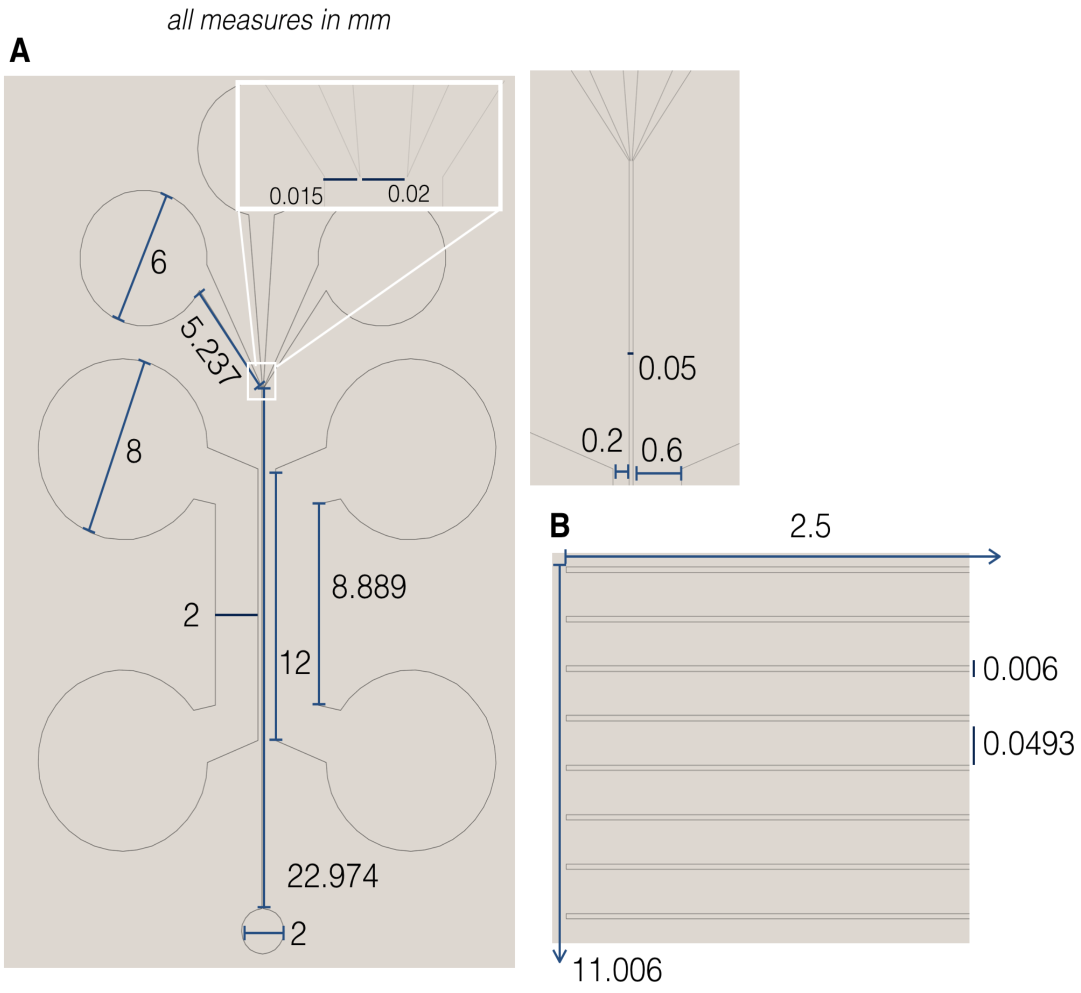


**Fig S1. Design of the photolithography masks. A.** Layout is based on (Taylor 2010). Changes have been introduced in many dimensions to increase the overall yield when cutting the synaptic compartment. The image shows the Mask containing the reservoirs as well as the perfusion channel, for second exposure. Height of the photoresist (SU-8-2050) for this step is 160µm. **B.** Mask containing the microgrooves, used for first exposure. Height of the photoresist (SU-8-2025) for this step is 20µm.

**Figure S2**

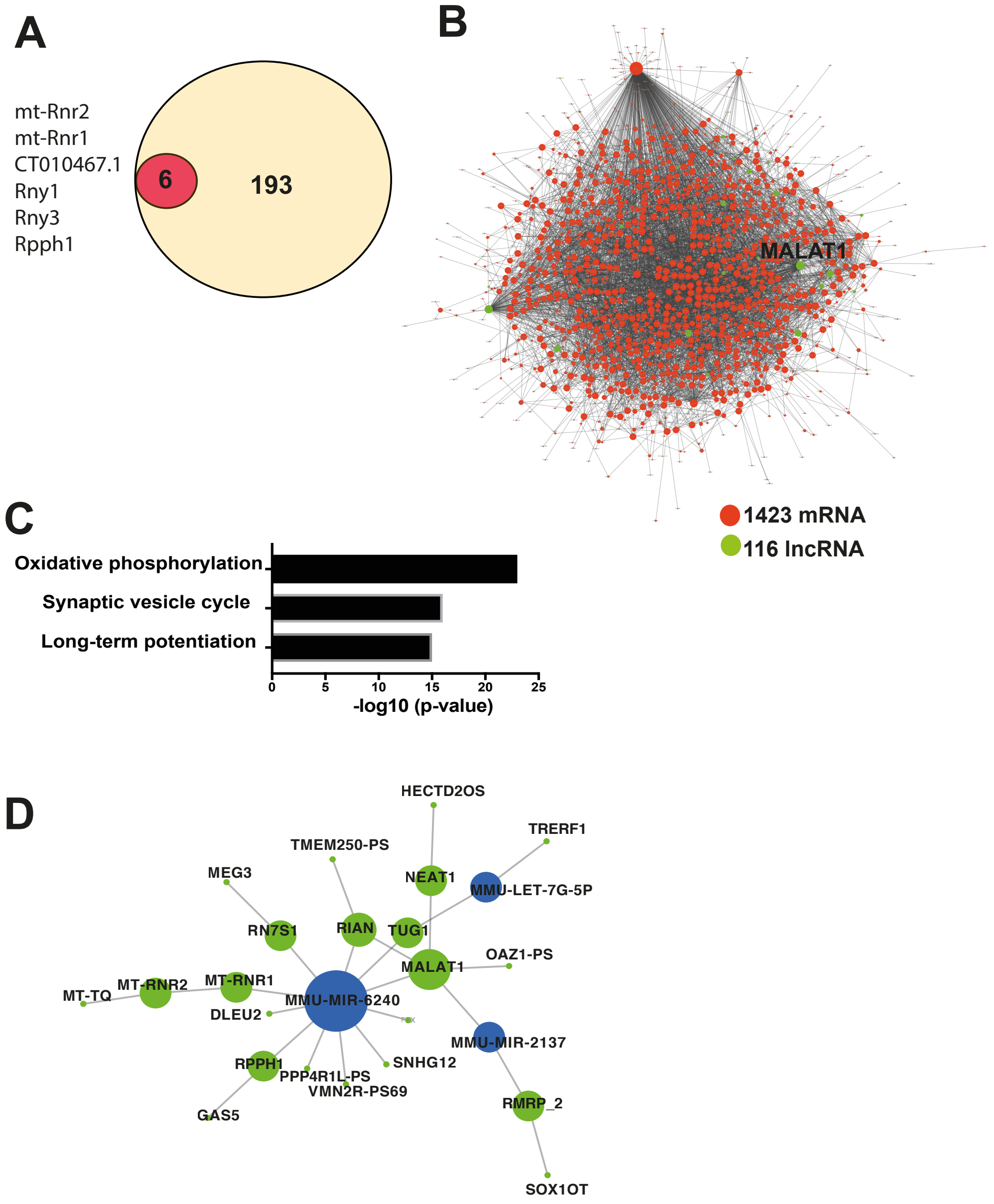


**Fig. S2. Network analysis of synaptic lncRNAs. A.** Venn diagram showing that the 6 annotated lncRNA detected in synaptosomes (red circle) are part of the 199 lncRNA detected in the microfluidic chambers. The 6 common lncRNAs are listed in the left panel. **B.** Out of 1460 mRNA detected in microfluidic chambers, 1423 could be fit into a network that also contains 116 of the 199 annotated lncRNA. Specifically Malat 1 appears as a hub lncRNA. **C.** Three main pathways represented by the synaptic mRNAs potentially controlled by the 116 lncRNAs. **D.** Network showing the interactions of synaptic lncRNA (green) and microRNA (blue) detected in microfluidic chambers. Note that the two well-studied lncRNAs Malat1 and Neat 1 are part of a network that controls microRNA-6240, let-7G-5p and microRNA-2137. Please note that panels B and D are available as searchable high resolution PDF files as additional supplemental data.
