## Supplementary material for "The Coding And Small-Non-Coding Hippocampal Synaptic RNAome": Table 1

**Table 1: Synaptic microRNAs detected in cognitive diseases**

| Synaptic microRNA[1] | Disease | Literature (PMID) |
|---|---|---|
| mmu-miR-7a-5p# | Depression | 32304043 |
| mmu-miR-125b-5p* | Alzheimer’s disease | 31240600 |
| mmu-miR-30d-5p# | Schizophrenia | 32304043 |
| mmu-miR-29a-3p# | Alzheimer’s disease; Schizophrenia | 31240600; 32304043 |
| mmu-miR-181a-5p# | Alzheimer’s disease | 31240600; 29121998; 32304043 |
| mmu-miR-30a-5p# | Schizophrenia | 32304043 |
| mmu-miR-125a-5p | Alzheimer’s disease | 31240600 |
| mmu-let-7d-5p* | Schizophrenia | 32304043 |
| mmu-miR-99b-5p# | Alzheimer’s disease | 29121998 |
| mmu-miR-26b-5p# | Schizophrenia | 32304043 |
| mmu-let-7e-5p* | Alzheimer’s disease | 29121998 |
| mmu-miR-29b-3p# | Schizophrenia | 32304043 |
| mmu-miR-30e-5p# | Schizophrenia | 32304043 |
| mmu-miR-24-3p* | Schizophrenia | 32304043 |
| mmu-miR-330-5p | Depression/Bipolar disease | 32304043 |
| mmu-miR-146a-5p# | Alzheimer’s disease | 31240600; 31437718 |
| mmu-miR-132-3p# | Alzheimer’s disease | 24014289; 31240600 |

**1:** ranked by expression level in synaptosomes (top to down)

*also detected in synaptic compartment of primary neurons grown in microfluidic chambers

#also detected in astrocytic exosomes
